# Bayes Factor for Linear Mixed Model in Genetic Association Studies

**DOI:** 10.1101/2024.05.28.596229

**Authors:** Yongtao Guan, Daniel Levy

## Abstract

Bayes factor has advantages over p-value as test statistics for association, particularly when comparing multiple non-nested alternative models. An efficient method to compute Bayes factor for linear mixed model in the context of genetic association studies is lacking. In this study, we transform the standard linear mixed model to a Bayesian linear regression by substituting the random effect with fixed effects, where the covariates of the fixed effects are eigenvectors of the genetic relatedness matrix and their respective prior effect sizes are proportional to the corresponding eigenvalues. Using conjugate normal inverse gamma priors on regression parameters, Bayes factors can be computed in a closed form. We demonstrate numerically the known relationship between Bayes factors and p-values for the linear mixed model. We then show that predictions based on the transformed Bayesian linear regression are identical to those of the best linear unbiased prediction (BLUP) of the standard linear mixed model. Our results provided a new perspective and derivation to a known connection between BLUP and Bayesian estimates. Methods described in this note are implemented in the software IDUL as two new functionalities: computing Bayes factors and residuals for the linear mixed model. IDUL and its source code are freely available at https://github.com/haplotype/idul.

## 1 Introduction

The linear mixed model (LMM) has emerged as an effective tool in controlling for population structure and relatedness in a study sample to reduce false positives in genetic association studies (Kang et al., 2008; Zhou and Stephens, 2012; Jiang et al.,2019). Consider a standard LMM in genetic association literature

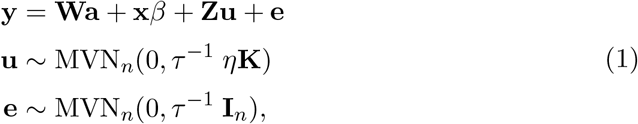

where **W** contains conventional covariates such as age and sex, including a column of 1; **x** contains genetic variant(s) to be tested for association; and **u** is the random effect with **Z** as its loading matrix and **K**, whose (*i, j*)-th entry is twice the kinship between samples *i* and *j*, as its covariance matrix (both **Z** and **K** are known). MVN_*n*_ denotes an n-dimensional multivariate normal distribution, **I**_*n*_ is n-dimensional identity matrix. In genetic association studies, the random effect **Zu** is a nuisance term that absorbs part of the phenotype **y** that is attributable to population stratification and relatedness. Here *η* is a hyperparameter, measuring the ratio between two dispersion terms (random effect **u** and random noise **e**).

In classical literature, *η* is known (Henderson, 1975). In GWAS applications, however, *η* is an important parameter to be estimated, either using maximimum likelihood (ML) or restriced maximum likelihood (REML) approaches (Zhou and Stephens, 2012; Guan and Levy, 2024a). Conditioning on *η*, estimates of 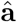 and 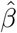 can be obtained and either the likelihood ratio test or Wald test can be used to compute a p-value to test against the following null model

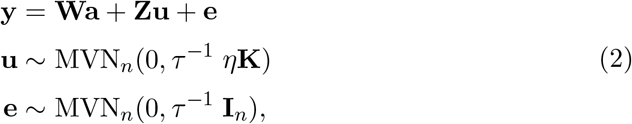

where the hyperparameter *η* was estimated from the alternative model (1). IDUL algorithm is an efficient and asymptotically exact algorithm to estimate *η* (Guan and Levy, 2024a), and IDUL software can compute likelihood ratio test p-value using ML estimate of *η* or Wald test p-value using REML estimate of *η*.

Bayes factors has advantages over p-values, particularly for comparing multiple non-nested alternative models (Kass and Raftery, 1995; Stephens and Balding, 2009). An efficient method to compute Bayes factors for LMM in the context of genetic association studies is missing. Our main objective here is to developing an efficient method to evaluate Bayes factors for the linear mixed model. In order to do that, we first transform the LMM (1) to a Bayesian linear regression with a specific prior, then use standard normal inverse gamma priors on regression parameters to express Bayes factors in a closed form, which can be evaluated efficiently.

The transformed Bayesian linear regression can be used for prediction, and we show that the Bayesian prediction is identical to that of best linear unbiased prediction (BLUP) (Robinson, 1991).

## 2 Results

### 2.1 Bayesian linear regression

With reference to Model (1), the genetic relatedness matrix can be computed as **G** = **ZKZ**^*t*^. Let the eigen decomposition of **G** as **G** = **QDQ**^*t*^, where **D** = *diag*(*d*_1_, …, *d*_*n*_) with *d*_1_ ≥ *d*_2_ ≥ *· · ·* ≥ *d*_*n*_ and **QQ**^*t*^ = **I**_*n*_. Model (1) can be rewritten as

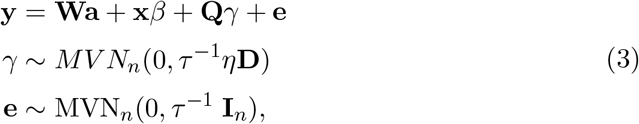

where the prior on *γ* has been specified. Model (3) is equivalent to model (1). To see this, *E*(**Zuu**^*t*^**Z**^*t*^) = *η***ZKZ**^*t*^ = *η***G** = *η***QDQ**^*t*^ = *E*(**Q***γγ*^*t*^**Q**^*t*^). Similarly, Model (2) can be rewritten as

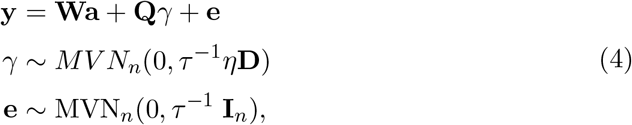

Note the priors on *γ* are the same between the null and the alternative models. The hyperparameter *η* can be estimated and plugged in; Its maximum likelihood estimate can be efficiently computed (c.f. Guan and Levy, 2024a).

### 2.2 Priors and Bayes factors

Let **X** = (**Q, W**). By specifying a Gamma prior on *τ*, we have the standard normal-inverse-gamma on a linear model (c.f Servin and Stephens, 2007; Zhou and Guan, 2018).

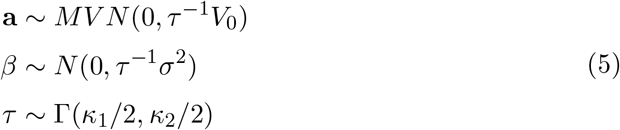

For Model (4), after integrating out **a**, *γ* and *τ* and letting *κ*_1_, *κ*_2_ → 0, we have

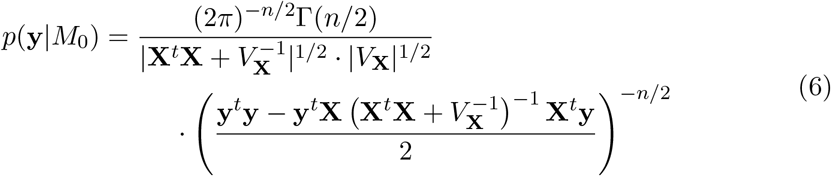

where *V*_**X**_ = diag 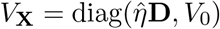. Similarly, for Model (3), we have

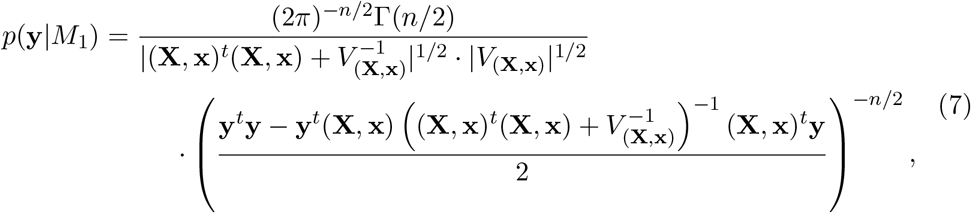

where *V*_(**X**,**x**)_ = diag 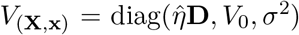. The Bayes factor *p*(**y**|*M*_1_)*/p*(**y**|*M*_0_), after letting *V*_0_ → **0**, is

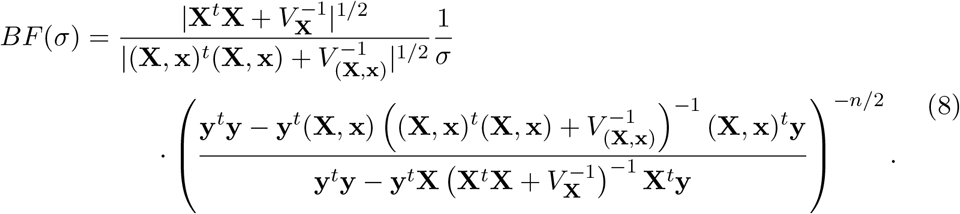

Note that here we used improper priors, but both the posterior and Bayes factor are proper (Servin and Stephens, 2007). The Bayes factor can be evaluated efficiently (Methods).

### 2.3 Numerical examples

Figure 1 shows comparisons between Bayes factors computed using Equation (8) and p-values from the likelihood ratio test calculated using IDUL software (Guan and Levy, 2024a), stratified according to minor allele frequencies (MAF) of the SNPs. The data, consisting 5757 whole genome sequenced samples with high level of relatedness and 79 protein assay phenotypes, were described previously (Guan and Levy, 2024b). We make the following observations: 1) In general, large Bayes factors correspond to small p-values. 2) Different Bayes factors might have the same p-values, and different p-values might have the same Bayes factors. 3) Left panel shows that, for SNPs with the same p-values, those having larger MAF tend to have larger Bayes factors than those having smaller MAF. This is because Bayes factors penalize SNPs of small MAF for their lower information contents through the prior effect size (Guan and Stephens, 2011). 4) Previously it has been demonstrated that asymptotically 2 log *BF* = *λχ*^2^ +log(1 −*λ*), where *λ* ∈ (0, 1) increases with the SNP MAF and is also a function of prior effect size (Zhou and Guan, 2018). Right panel reflects the mathematical insight between Bayes factors and their corresponding *χ*^2^-values.

**Figure 1.**
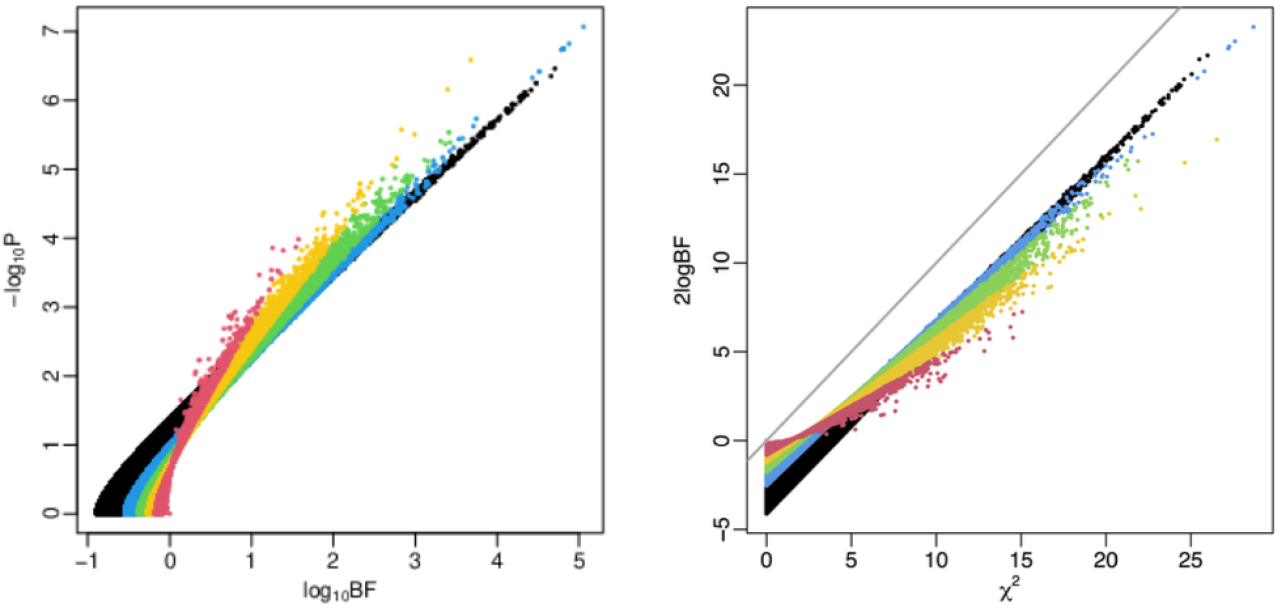
Comparison between p-values and Bayes factors for linear mixed model. The left panel shows log_10_ *BF* versus −log_10_ *P*. The right panel shows *χ*^2^ versus 2 log *BF*, where *χ*^2^ values are quantiles of the p-values. The gray line is *x* = *y*. The color is based on MAF *f*, with black for *f* ∈ (0.05, 0.5], blue for *f* ∈ (0.02, 0.05], green for *f* ∈ (0.01, 0.02], orange for *f* ∈ (0.005, 0.01], and red for *f* ∈ (0, 0.005].

### 2.4 Bayesian prediction and BLUP

From model (4) we estimate parameters of 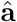 and 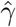 by 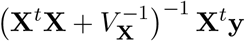 (Methods). Note both are column vectors. For a new sample, we have a row vector **w** for covariates and a row vector **g** for genetic relatedness between the new sample with *n* existing samples. Then the Bayesian prediction for the new sample is

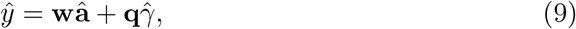

where **q** = **gG**^−1^**Q** is the least square prediction (based on genetic relatedness) of the new sample onto **Q** (Methods). For an existing sample *j*, **q** is the *j*-th row of the matrix **Q**.

We now show that the best linear unbiased prediction (BLUP) is identical to Bayesian prediction. From the Frequentist point of view, **u** is a random vector instead of a fixed parameter, it can only be *predicted* instead of *estimated*. The Bayesian interpretation of BLUP is well known (Gianola and Fernando, 1986; Robinson, 1991). Here we offer a new derivation that is perhaps more natural from the perspective of genetic association studies. For BLUP one first estimates 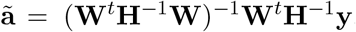, where 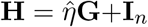, and then estimate 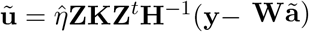. Note both are column vectors. Thus for the new sample, BLUP prediction is

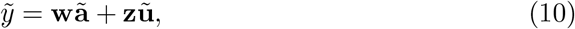

where **z** = **gG**^−1^**Z** is the least square prediction (based on genetic relatedness) of the new sample onto **Z** (Methods). For an existing sample *j*, **z** is the *j*-th row of the matrix **Z**.

To show Bayesian prediction 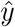 in (9) is identical to BLUP 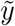 in (10), it is sufficient to show that 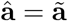 and 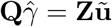, which we defer to Methods. Therefore, BLUP is identical to Bayesian prediction under a specific prior. For the existing samples, we can compute residuals using either Bayesian estimates or BLUP as following:

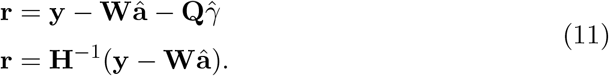

Residuals are useful in removing spurious correlations due to population structure and (cryptic) relatedness for subsequent analysis, such as studying gene-gene interaction and gene networks using gene expression values.

## 3 Discussion

It is a common practice to control for top principal components (PCs, the columns of the **Q** matrix) in genetic association study to control for population stratification (Price et al., 2006). It is self evident that if a putative causal SNP is correlated with both phenotype and top PCs, this practice of controlling for top PCs will reduce power. Our derivation suggests that when controlling for top PCs, one should put prior on their effect sizes 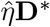, where **D**^∗^ is the a submatrix of **D** corresponding to the top PCs. Intuitively, the prior will keep some variations of PCs in the phenotypes for the putative causal SNPs, and a priori, the eigenvalues are the right amount of variation to be controlled for.

In classical literature dealing with linear mixed model (Henderson, 1975; Robinson, 1991), the covariance structure of the error term is a more general **R** instead of our diagonal **I**_*n*_. We can left multiplying 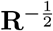 (where **R**^−^ is a generalized inverse) to convert the classical form to the form in Equation (1). In fact, we can rotate the system further and compute the (empirical) Bayes factor based on a rotated simple linear regression. Multiplying both side of Equation (4) by **Q**^*t*^ we get 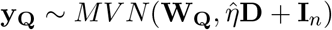, where **y**_**Q**_ = **Q**^*t*^**y** and **W**_**Q**_ = **Q**^*t*^**W**. Let Λ = *η***D** + **I**_*n*_ and scale **y**_**Q**_ and **W**_**Q**_ by left multiplying 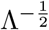, we get

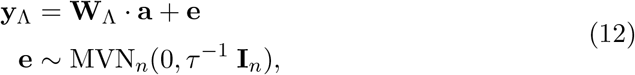

where 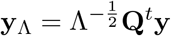 and 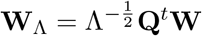. Similarly Equation (3) becomes

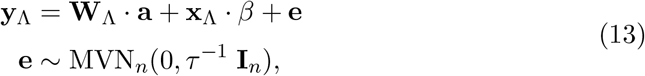

where 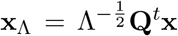. Then using priors specified in (5), we can compute Bayes factors (Servin and Stephens, 2007). This rotation approach was briefly mentioned by (Wen, 2015). Bayes factor from this approach in equivalent to Equation (8), but this approach obscures insights on Bayesian prediction and its equivalence to BLUP. We took an empirical Bayesian approach and plugged in ML estimate of the hyperparameter 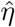. Of course, we can take the full Bayesian approach by coming up with a prior for *η*. One sensible choice is to define *h* = *η/*(1 + *η*) and assign a uniform prior Unif(0, 1) on *h*. Then *η* has a probability density function *f* (*η*) = 1*/*(1 + *η*)^2^, which is heavy tailed (Guan and Stephens, 2011). This full Bayesian approach is advantageous in principle, as it accounts for variation of *η*, but the computation is difficult, as this prior cannot be integrated out analytically, and either sampling or numerical integration are required for computation. For example, Wen (2015) took a full Bayesian approach assigning priors on *η* and *τ*, and elected to compute approximate Bayes factors for linear mixed model. On the other hand, by plugging in ML estimate 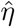, the subsequent computation can be done analytically. And this empirical Bayes approach is considered approximation to the full Bayesian treatment (Carlin and Louis, 2002).

Finally, Bayes factor is more desirable than p-values in applications that require comparing multiple non-nested alternative models. One such example is comparing genotype effect, paternal effect, and maternal effect, and paternal-maternal joint effects of the same SNP. More details can be found in our manuscript on parent-of-origin effect eQTL (Guan et al., 2025). Methods developed here are implemented in a software package that is freely available at https://github.com/haplotype/idul.

## 4 Methods

### 4.1 Computation

Bayes factor (8) can be evaluated efficiently. This computation was documented previously (Guan and Levy, 2024b), and we reproduced it here for context clarity. As 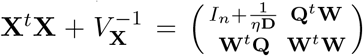 and denote 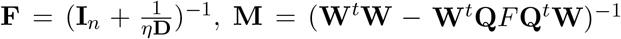, we compute its determinant using the identity 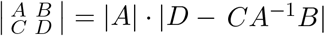 to get

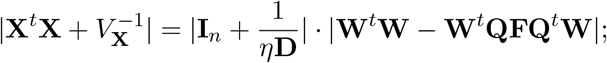

Using the identities 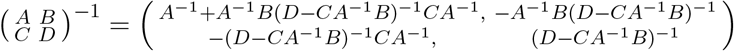,we get

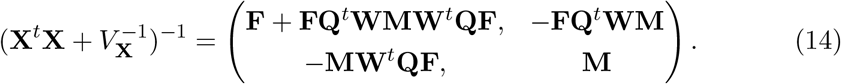

Note that for large matrices the computation only involves an inexpensive matrix multiplying vector, without expensive matrix multiplication and matrix inversion. For small matrices such **W**^*t*^**W** and *M*, the multiplication and inversion are inexpensive. The only expensive calculation is the eigendecomposition to obtain Q and D, which only needs to be done once. Similarly, both 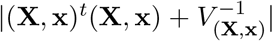 and 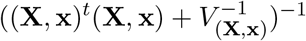 can be efficiently computed. Finally,

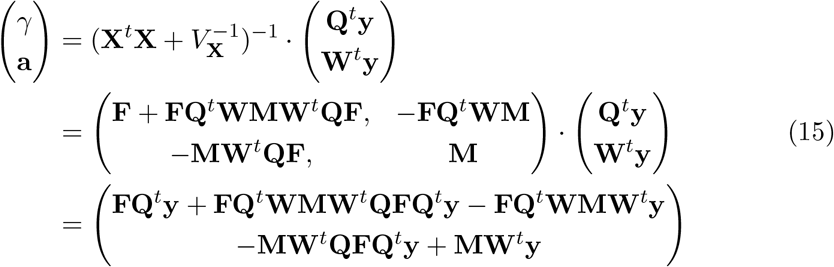

### 4.2 Least square projection

Let **Q**_*j*_ be the *j*-th column of **Q**, and consider model **Q**_*j*_ = **G***β*_*j*_ + *ϵ*_*j*_. The least square estimates of *β*_*j*_ = **G**^−1^**Q**_*j*_. For a new observation **g**, the prediction on the **Q**_*j*_ is **gG**^−1^**Q**_*j*_. Since this is true for each *j*, we have the least square prediction onto **Q** based on genetic relatedness is **gG**^−1^**Q**. The same arguments apply to **Z** as well.

### 4.3 Equivalence

With the notations defined in the main text, we aim to show 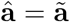 and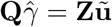.

With reference to (15) and the fact that **QFQ**^*t*^ = **I**_*n*_ − **H**^−1^, we have

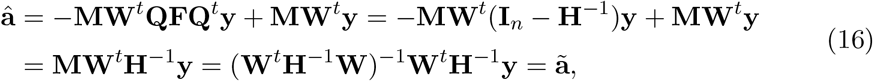

and

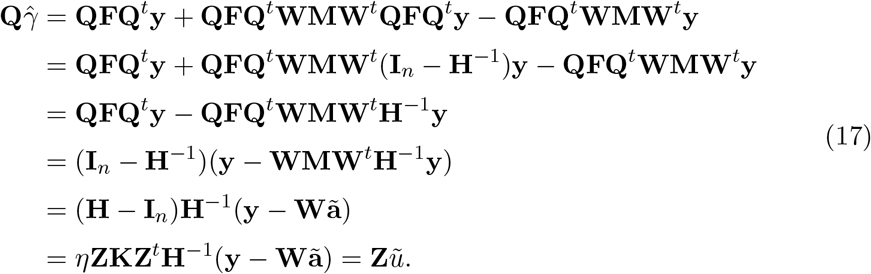

